# The Cholesteryl Ester Transfer Protein (CETP) raises Cholesterol Levels in the Brain and affects Presenilin-mediated Gene Regulation

**DOI:** 10.1101/2020.11.24.395186

**Authors:** Felix Oestereich, Noosha Yousefpour, Ethan Yang, Alfredo Ribeiro-da-Silva, Pierre Chaurand, Lisa Marie Munter

## Abstract

The cholesteryl ester transfer protein (CETP) is a lipid transfer protein responsible for the exchange of cholesteryl esters and triglycerides between lipoproteins. Decreased CETP activity is associated with longevity, cardiovascular health, and maintenance of good cognitive performance. Interestingly, mice lack the CETP-encoding gene and have very low levels of low-density lipoprotein (LDL) particles compared to humans. To understand how CETP activity affects the brain, we utilised CETP transgenic (CETPtg) mice showing elevated LDL levels on a high cholesterol diet inducing CETP expression. We found that CETPtg mice had up to 25% higher cholesterol levels in the brain. Using a microarray on astrocyte-derived mRNA, we found that this cholesterol increase is likely not due to astrocytic-dependent *de novo* synthesis of cholesterol. Rather, several genes linked to Alzheimer’s disease were altered in CETPtg mice. Most interestingly, we found activation of the G-protein coupled receptor EP4 and γ-secretase as upstream regulators of these transcriptional changes. Further *in vitro* studies showed that CETP expression was sufficient to activate γ-secretase activity. The data suggest that CETP activity affects brain’s health through modulating cholesterol levels and Alzheimer’s-related pathways. Therefore, CETPtg mice constitute a valuable research tool to investigate the impact of the cholesterol metabolism on brain functions.

## Introduction

Cholesterol is a major constituent of biomembranes and precursor for various hormones. In most tissues, the cholesterol concentration is about 2 mg/g tissue, however, it reaches 15-20 mg/g in tissue of the central nervous system (CNS) (1). Thus, the brain contains 25% of the total body cholesterol, suggesting a special need of the brain for cholesterol (2). In the blood, dietary cholesterol is transported by very-low density lipoprotein (VLDL) or low-density lipoprotein (LDL) particles that are secreted by the liver to deliver cholesterol to extrahepatic tissues (3). Reverse cholesterol transport from the periphery back to the liver occurs via high-density lipoprotein (HDL) particles (4). However, the brain seems to be excluded from these distribution cycles since neither VLDL or LDL particles cross the blood-brain barrier (2,5–7). In the CNS, astrocytes are the cell type primarily involved in lipid synthesis and secrete HDL-like lipoprotein particles that contain predominantly apolipoprotein E (ApoE) as the apolipoprotein (8). Such particles are taken up by neurons through members of the LDL-receptor family that recognise ApoE including the LDL-receptor related protein 1 (LRP1) (9).

The cholesteryl ester transfer protein (CETP) is a lipid transfer protein that facilitates the exchange of cholesteryl esters in HDL for triglyceride in VLDL and LDL (10,11). The net result of this transfer activity is increased cholesterol content in pro-atherogenic LDL particles and decreased cholesterol levels in anti-atherogenic HDL particles (12). Studies investigating the genetic predisposition of “super-agers” or “centenarians” with well-maintained health and cognitive performance, revealed that polymorphisms that impair CETP’s activity associate with longevity, cardiovascular health, and good cognitive performance (13–15). Based on these findings, several studies investigated whether CETP polymorphisms could decrease the risk for Alzheimer’s disease, an aging-associated neurodegenerative disease. Indeed, protective effects of CETP polymorphisms at *early* Alzheimer’s disease stages were reported, particularly in carriers of the strongest genetic risk factor, the ∊4 allele of the apolipoprotein E (ApoE4) (16–19). ApoE is the predominant lipoprotein of the brain, in contrast to the blood where there are several apolipoprotein-defined lipoprotein families (20). Those epidemiological findings indicate that CETP activity may impact on cognitive performance and brain functions, however, the underlying molecular mechanisms remain unclear.

While CETP is predominantly expressed in the liver and secreted to the blood, it is expressed in astrocytes as well (21). However, its function in the CNS remains elusive. Considering the important effect of CETP on systemic cholesterol levels, we hypothesised that CETP may also contribute to the alterations of the brain’s cholesterol levels. It is important to note that mice naturally lack CETP and therefore they have considerably less LDL compared to humans (22). To gain insight on how CETP may impact on cognitive performance in humans, we used a well-established CETPtg mouse model expressing the human CETP gene under its natural promoter (CETPtg) that is frequently used in the cardiovascular research field (23). The promoter contains a cholesterol responsive element that induces CETP gene expression in response to dietary lipids. Therefore, CETP expression in CETPtg mice leads to increased LDL levels and could thus be regarded as a mouse model with a humanised (normolipidemic) lipoprotein profile (24). We herein characterised the effects of CETP expression on molecular changes in the brain in CETPtg mice. We observed higher cholesterol levels in the brains of CETPtg as compared to wild type (wt) mice. Transcriptome profiling of astrocytes indicated decreased cholesterol synthesis, and modulation of several genes linked to Alzheimer’s disease including an overall activation of presenilin-mediated signaling.

## Methods

All experiments were conducted in accordance with McGill University environmental health and safety regulations (EHS) as well as the Canadian biosafety standards and guidelines.

### Cell culture

HEK293T cells were cultivated in 1:1 Dulbecco’s modified Eagle medium (DMEM) supplemented with 0.584 g/l L-glutamine and 0.11 g/l sodium pyruvate (Wisent), and 10% FCS (Wisent), at 37°C and 5% CO_2_. For transient transfections, 1.5 × 10^5^ cells per well (12-well plates) were seeded 24 h before transfection. Cells were transiently transfected with 1 μg DNA in total and 2 μl polyethyleneimine (PEI) per well. 36 hours after transfection, cell culture supernatant was collected, and cells were lysed with TNE-lysis buffer (50 mM Tris, pH 7.4, 150 mM NaCl, 2 mM EDTA, 1% NP40, and complete protease inhibitors, Roche) and prepared for SDS-polyacrylamide gel electrophoresis (SDS-PAGE).

### Western blot analysis of mouse tissue samples

Fresh frozen liver or brain samples (approximately 100 mg) were lysed in 5x volume of lysis buffer (150 mM NaCl, 10% Glycerol, 2 mM EDTA, 0.5% NP-40, 0.1% sodium-deoxycholate, 20 mM HEPES, 1x complete protease-inhibitor cocktail (Roche), pH 7.4) using lysing-matrix D at 6000 rpm for 40 seconds. The lysates were further diluted 1:5 in lysis buffer. For western-blot analysis, liver samples were prepared for SDS-polyacrylamide gel electrophoresis (SDS-PAGE) and loaded on either 10% or 15% SDS-polyacrylamide gels. The following primary antibodies were used: 22C11 (Millipore), rabbit-anti-GAPDH (14C10, Cell Signaling), TP2 (kind gift of the Ottawa Heart Institute), anti-TREM2 (Mab1729 R&D systems) and anti-ABCA7 (polyclonal, Thermo Fisher). Horseradish peroxidase (HRP)-coupled secondary antibodies directed against mouse or rabbit IgG were purchased from Promega. Chemiluminescence images were acquired using the ImageQuant LAS 500 system (GE Healthcare).

### Quantitative real time PCR (RT-qPCR)

mRNA was isolated from mouse tissue using the Macherey & Nagel mRNA-isolation kit in combination with lysing matrix D. Briefly, 25-50 μg of fresh frozen tissue were lysed in 450 μL RNA preparation buffer (with β-mercaptoethanol) in lysing matrix D tubes using a magna lyser (6000 rpm 2x 30 seconds) according to manufacturer’s instructions. The RNA concentration was adjusted to 100 pg/mL and 500 ng of RNA were transcribed into cDNA using the high-capacity cDNA reverse-transcription kit (Applied Biosystems) according to manufacturer’s instructions. RT-qPCR was performed using the SsoAdvanced SYBR green supermix (Biorad) according to manufacturer’s instructions on a Biorad CFX384Touch cycler. All primers were ordered from integrated DNA technologies. Primers used were: CETP forward: CAGATCAGCCACTTGTCCAT, CETP reverse: CAGCTGTGTGTTGATCTGGA, ABCA7 forward: TTCTCAGTCCCTCGTCACCCAT, ABCA7 reverse: GCTCTTGTCTGAGGTTCCTCGT, TNFα forward: GGTGCCTATGTCTCAGCCTCTT, TNFα reverse: GCCATAGAACTGATGAGAGGGAG, IL1β forward: TGGACCTTCCAGGATGAGGACA IL1β reverse: GTTCATCTCGGAGCCTGTAGT, TLR4 forward: AGCTTCTCCAATTTTTCAGAACTTC, TLR4 reverse: TGAGAGGTGGTGTAAGCCATGC, TREM2 forward: ACAGCACCTCCAGGAATCAAG, TREM2 reverse: AACTTGCTCAGGAGAACGCA, IL6 forward: CCTCTGGTCTTCTGGAGTACC, IL6 reverse: ACTCCTTCTGTGACTCCAGC, HES1 forward: p21 forward: GCCTTAGCCCTCACTCTGTG p21 reverse: AGCTGGCCTTAGAGGTGACA, HES1 forward: CGGAATCCCCTGTCTACCTC, HES1 reverse: AATGCCGGGAGCTATCTTTCT. The following primers were used as reference genes: HPRT forward: CCAGTTTCACTAATGACACAAACG, HPRT reverse: CTGGTGAAAAGGACCTCTCGAAG, PSMC4 forward: CCGCTTACACACTTCGAGCTGT, PSMC4 reverse: GTGATGTGCCACAGCCTTTGCT, GAPDH forward: CATCACTGCCACCCAGAAGACTG, GAPDH reverse: ATGCCAGTGAGCTTCCCGTTCAG, Actin-β forward: CATTGCTGACAGGATGCAGAAGG, Actin-β reverse: TGCTGGAAGGTGGACAGTGAGG. Primer efficiency for all primers was determined to be between 90-110%. For normalization of gene expression, the four genes ACT, GAPDH, HPRT and PSMC4 were used as reference genes. RT-qPCR was analysed using the CFX manager software (Biorad).

### Imaging mass spectrometry (IMS)

Sample preparation: The fresh frozen brain samples were sectioned sagittally at 14 μm thickness and the frozen brain homogenates at 20 μm thickness with a Leica CM3050 cryostat at −20°C (Leica Microsystems GmbH, Wentzler, Germany). All brain specimens were cut at approximately the same Bregma in order to clearly delineate the hippocampus. Brain homogenates were prepared according to published protocols (25) and were used to normalise data across experiments. For each technical replicate, one tissue section of each condition was thaw-mounted in a 2 × 2 pattern on a 25 × 75 mm indium-tin-oxide (ITO) coated microscope slide (Delta Technologies, Loveland, CO), along with two sections of frozen brain homogenate on the left and right of the grid. After desiccation in a vacuum pump desiccator for ≤ 1 hour, a 23 ± 2 nm silver layer was deposited onto the sections using a Cressington 308R sputter coater (Ted Pella Inc, Redding, CA) as per the protocol detailed in Dufresne et al 2013 (26). The argon partial pressure was set at 0.02 mbar and the current at 80 mA. Data acquisition: IMS data were acquired at 50 μm spatial resolution and 100 shots per raster position with a “small” laser setting using a “matrix-assisted laser desorption/ionization-time of flight (MALDI-TOF/TOF) ultrafleXtreme mass spectrometer (Bruker Daltonics, Billerica, MA) equipped with a SmartBeam-II Nd:YAG/355-nm laser operating at a repetition rate of 1 kHz using flexImaging 4.1 software (Bruker Daltonics, Billerica, MA). All instrumental parameters (source voltages, laser energy, delayed extraction parameters, etc.) were optimised for maximum signal-to-noise ratio within the 100-1100 *m/z* range in the reflectron geometry, with the acceleration voltage set to 25 kV. Two 400-pixel squares were also acquired from each brain tissue homogenate section at the same spatial resolution. Data Analysis: Raw IMS data were first internally calibrated with the silver isotopic peaks using the flexAnalysis Batch Process software (Bruker Daltonics, Billerica, CA) to obtain a ~5 ppm mass accuracy. Next, IMS data from the hippocampal and whole brain regions of interests (ROIs) were exported into the common imzML format using flexImaging 4.1 (27). Using an in-house code based on the Cardinal package (x1.6.0) in R (x3.2.5), the mean area and standard deviation of the two cholesterol signals (*m/z* 493.26 and *m/z* 495.26, corresponding to the [M+^107^Ag]^+^ and [M+^109^Ag]^+^ molecular ions, respectively) were calculated for the ROIs after independent TIC normalization (28). The same code was used to obtain the mean of the summed areas of the ten most abundant signals in the homogenate squares. This value acted as the correction factor to correct for variations in signal intensity across all experiments. The final cholesterol intensity reported is the mean across the three technical triplicates for one group normalised against the correction factor. Unless otherwise noted, all solvent and material were purchased from Thermo Fisher Scientific (Ottawa, ON). The silver target 3N5 (99.95% purity) used for tissue sputter-coating was purchased from ESPI Metals (Ashland, OR).

### Filipin staining and Immunohistochemistry

Fresh-frozen brains were cut on the sagittal plane at 25 μm thickness using a cryostat (Leica, Germany). Sections were then collected and fixed in 4% paraformaldehyde at 4°C for two hours. Filipin III (stock powder) was dissolved at a concentration of 10 mg/ml in dimethylformamide (DMF) and diluted 100-fold with 10 mM phosphate buffered saline (PBS). For Filipin staining, tissues were washed in PBS and incubated in 0.01 mg/ml Filipin complex solution (Sigma Aldrich) at room temperature for two hours. After washing with PBS, brain sections were mounted on a microscope slide and imaged with Zeiss AxioImager M2 Imaging microscope with the Zeiss ZenPro software v.2.3 (Zeiss Canada). For Immunohistochemistry, brain sections were permeabilized with 0.2% Triton-X in PBS (PBST) and blocked for 1 hour at room temperature in 10% normal donkey or goat serum. Sections were incubated in a cocktail of primary antibodies composed of mouse anti-GFAP (Cell signaling, cat # 3670, 1:1000), and rabbit anti-C1q (Abcam, cat # ab182451, 1:400) prepared in 5% blocking solution for 12 h at 4°C. Primary antibody labelling was detected using species-specific secondary antibodies conjugated to Alexa 488, and Alexa 568 (Invitrogen, 1:800, incubated at room temperature for 2 hours). Sections were mounted on gelatin subbed slides and coverslipped using Prolong Gold Antifade mounting medium (Invitrogen) and Zeiss cover slips. Sections were imaged using Zeiss LSM 800 confocal microscope. For image analysis, Filipin and C1q fluorescence intensity levels were quantified by the average intensity of staining in ImageJ (NIH) using images captured by 20x objective (C1q) and 40x objective (filipin). Specifically for filipin staining, all images were thresholded to an equal value that was determined empirically and only fluorescent intensities corresponding to the cell membranes were quantified. All values were normalized to the background fluorescence of the corresponding image.

### Mouse housing

The CETPtg mouse strain B6.CBA-Tg(CETP)5203Tall/J (Jackson strain no.: 003904) (23) were housed according to the McGill University standard operating procedure mouse breeding colony management #608. All procedures were approved by McGill’s Animal Care Committee and were performed in accordance with the ARRIVE guidelines (Animal Research: Reporting in Vivo Experiments). Mice were bred heterozygous and non-transgenic littermates were used as controls. Mice of both sexes were used in analyses at about equal ratios. All mouse diets were purchased from Envigo. The diets used in this study were: low fat control diet (TD.08485), low fat diet enriched with 1% cholesterol (TD.140215) and a diet containing 21% fatty acids (FA) and 1% cholesterol (TD.95286). The FA composition was 65% saturated FA (SFA), 31% monounsaturated FA (MUFA), and 4% polyunsaturated FA (PUFA). Animals of both sexes were assigned randomly to treatment groups.

### Mouse genotyping

Genotyping was performed by Transnetyx genotyping using real-time PCR from ear punch tissue. Ear punches were lysed in at 56 °C overnight. Primers used for the transgene were: forward: GAATGTCTCAGAGGACCTCCC, reverse: CTTGAACTCGTCTCCCATCAG. Primers for internal controls were: Forward: CTAGGCCACAGAATTGAAAGATCT, reverse: GTAGTGGAAATTCTAGCATCATCC.

### Plasma lipid analysis

The lipid analysis of mouse plasma samples was performed using the COBAS Integra 400 Plus (ROCHE) and the following kits: COBAS INTEGRA CHOL 2, COBAS INTEGRA HDL-C gen3, and COBAS Integra TRIG GPO 250, respectively. The levels of LDL-C were calculated using the Friedewald formula: [total cholesterol] – [HDL-C] – [TG/2.2].

### CETP activity assay

CETP activity was measured using the Roar biomedical Inc. fluorescent CETP activity assay. Here, 5 μL of cell culture supernatant was incubated with 0.3 μL donor and 0.3 μL acceptor molecules in 30 μL reaction volume. The reaction mix was incubated for 3 hours at 37 °C in a water bath and the fluorescence (λex 465/λem 535) was measured.

### Astrocyte purification

Astrocytes were purified using the Anti-GLAST (ACSA-1) MicroBead Kit (Miltenyi biotec). Briefly, whole mouse brains were dissociated using a miltenyi gentleMACS Octo dissociator with Heaters, and GLAST positive astrocytes were isolated using anti-GLAST (ACSA-1) antibody magnetic beads according to manufacturer’s instructions.

### Flow cytometry

Purified astrocytes were EtOH fixed and stained with a Cy3 labelled anti-GFAP antibody (1:1000, Sigma). The samples were run on a BD LSRFortessa flow cytometer and the GFAP-Cy3 emission was detected using a 561 nm laser for excitation. The detector channel used was 586/15 nm. BD FACSDIVA 8.0.1. was used for analysis.

### Astrocyte microarray

RNA from GLAST-positive astrocytes was isolated using the Macherey & Nagel mRNA isolation kit. The Affymetrix clariom-S nano microarray was performed at the Genomecenter Quebec according to manufacturer’s instructions. The initial microarray dataset was analysed using Transcriptome Analysis Software (Affymetrix). Upstream regulator and pathway analyses were performed using Ingenuity Pathway Analysis (IPA). GEO accession number: GSE111242.

### Statistical analysis

Statistical analysis was performed using the Graphpad Prism 7 and 8 software.

Analyses include only parametric tests: students T-test, and two-way ANOVA followed by Bonferroni corrections and Tukey’s multiple comparison tests. All p values, statistical tests, N values and the experimental unites employed are indicated in figure legends.

## Results

### Dietary cholesterol intake induces CETP expression

CETPtg animals have been widely used in cardiovascular research (23). However, it remained unclear if fatty acids further induce CETP expression in addition to dietary cholesterol. To identify the ideal diet for inducing CETP expression in the CETPtg model, we compared a diet enriched with 1% (w/w) cholesterol to a diet containing 1% cholesterol plus 21% (w/w) fatty acids (cholesterol/FA) for their effects on CETP expression in CETPtg mice. Wt and CETPtg mice received diets for one month starting at 2 months of age (**Figure 1A**). As expected, CETPtg, but not wt mice showed CETP activity, confirming that there is no compensatory mechanism for the lack of CETP in wt mice. CETP activity was increased 2-fold in animals that were on a diet enriched in cholesterol or cholesterol/FA as compared to mice on standard diet (**Figure 1B**). Likewise, both high fat diets induced a 2-fold increase of circulating CETP protein levels in mouse plasma, as determined by western blot (**Figure 1D, E**). Further, we quantified CETP mRNA levels in liver by RT-qPCR and found that the diet supplemented only with cholesterol induced a bigger increase of CETP mRNA levels (8.8-fold) as compared to the high cholesterol/FA diet (7-fold) (**Figure 1C**). When assessing the lipid profile in plasma, we found that CETPtg mice had lower HDL levels on standard and high cholesterol diets, an effect that was missing in mice receiving the cholesterol/FA diet (**Figure 1F**). LDL cholesterol levels were significantly elevated in CETPtg animals fed with a cholesterol-enriched diet, which was not observed in animals fed with a cholesterol/FA diet, although both diets led to a similar increase in CETP activity and protein levels (**Figure 1G**). It is important to note that those LDL levels of approximately 1.2 mmol/L observed in CETPtg mice are still relatively low considering that human LDL levels < 3 mmol/L are still being considered healthy levels. Total cholesterol was not significantly affected by the diets (**Figure 1H**). However, we found a trend towards decreased levels of triglycerides in animals fed with the cholesterol diet independent of the genotype (**Figure 1I**). Finally, we analysed the net weight gain of mice during the 4-week diet period. In contrast to the cholesterol/FA diet, mice on the cholesterol diet did not show an additional weight gain as compared to standard diet (**Figure 1J**). Together, high CETP expression and enhanced activity are achieved with both diets. However, the blood lipoprotein profile only changed towards a more human-like profile, i.e. increased LDL levels, in mice receiving the cholesterol-only diet. In addition, since cholesterol-enriched food did not impact the weight of mice, this diet has the advantage that potentially confounding factors such as obesity can be excluded. Thus, we chose to use the 1% cholesterol diet for further experiments.

**Figure 1:**
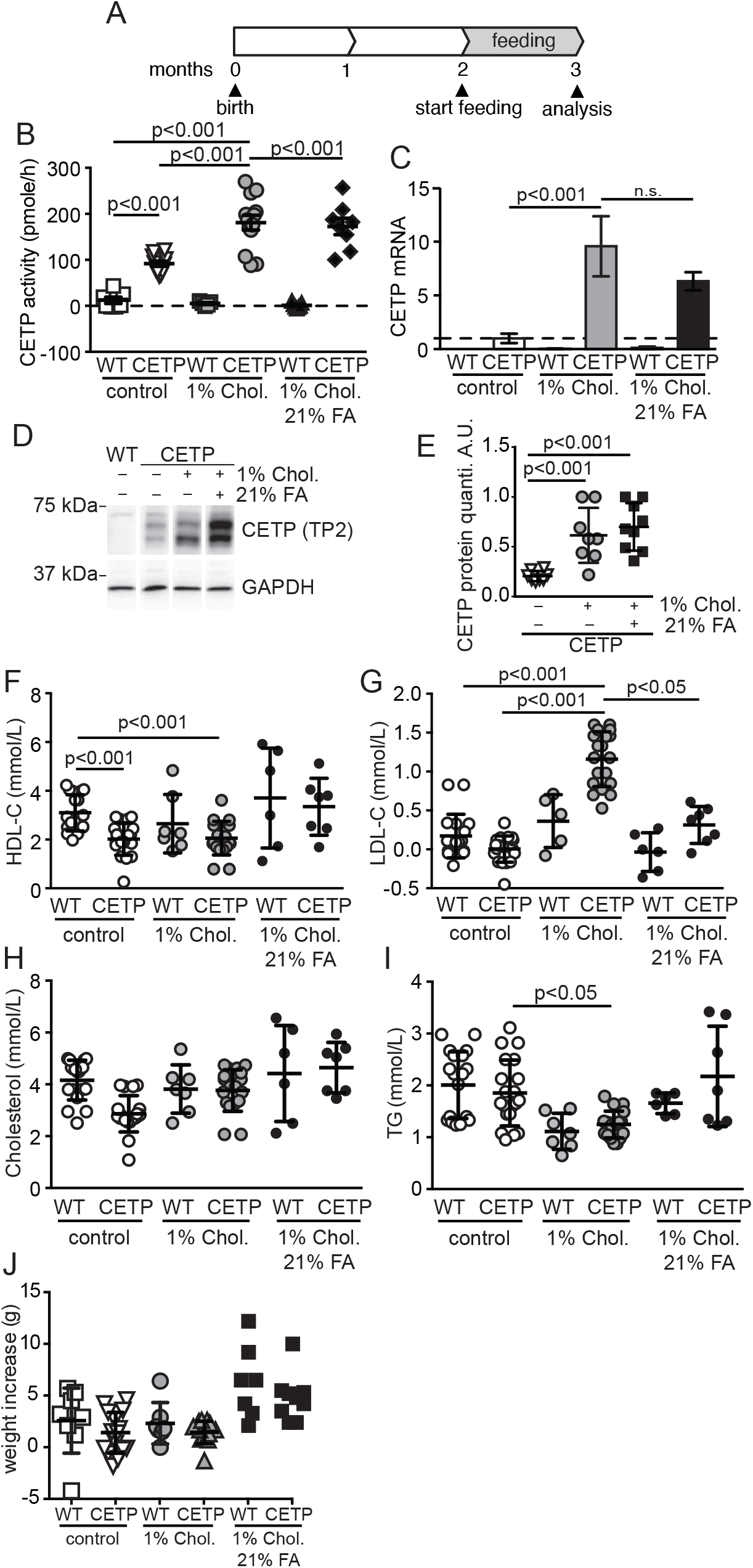
Dietary cholesterol intake induces CETP expression: **A**: Feeding schedule & study design. Wt and CETPtg animals were fed for 1 month starting at the age of 2 months. Biochemical analyses were performed after 3 months of age. **B:** CETP activity of CETPtg or wt animals was measured from 1 μL plasma using the fluorescence-based CETP activity assay (Roar biomedical). n=6-14, mean ± SEM; 2-way ANOVA, Tukey’s multiple comparison. **C:** Relative normalised CETP expression. RT-qPCR of liver samples at the age of 5 months. n=5-8, mean ± SEM; 2-way ANOVA, Tukey’s multiple comparison. **D:** CETP western blot from liver lysates. Liver lysates were separated on 10% SDS-PA gels. CETP was detected using the TP2 monoclonal antibody. **E:** Quantification of CETP western blots as shown in **D:** n=8, mean ± SEM; Students T-test. **F-I:** Plasma lipoprotein analysis: **F:** HDL-C, **G**: LDL-C, **H:** total cholesterol **I:** and triglycerides from mouse plasma samples. Plasma samples were analysed on a COBAS Integra 400 Plus (ROCHE) analyser using the following kits: COBAS INTEGRA CHOL 2, COBAS INTEGRA HDL-C gen3, and COBAS Integra TRIG GPO 250, respectively. The levels of LDL-C were calculated using the Friedewald formula: [total cholesterol] – [HDL-C] – [TG/2.2]. n=6-14. mean ± SEM; 2-way ANOVA, Tukey’s multiple comparison. **J:** Mouse weight increase: Net weight increase of wt and CETPtg mice during the feeding period.

### CETP promotes TREM2 expression in the liver

To ultimately study the chronic effect of CETP expression on the brain, we expanded the diet period to 3 months (**Figure 2A**). First, we analyzed the effect of CETP on plasma LDL levels and transcriptional changes in the liver. Dietary cholesterol is known to decrease cholesterol synthesis and transcription of genes involved in cholesterol synthesis (the rate-limiting enzyme 3-hydroxy-3-methylglutaryl-coenzyme A reductase, HMGCR) and cholesterol uptake (LDL-receptor, LDLR, and the lipoprotein-receptor-related protein-1, LRP1) through regulation of the sterol regulatory element-binding protein-2 (SREBP-2) (29). Indeed, HMGCR, LDLR, and LRP1 mRNA levels were decreased on high cholesterol diet in wt and CETPtg mice as compared to wt mice on a standard diet at the age of 5 months (**Figure 2 B-D**) (30). In addition, CETPtg mice on a standard diet showed lower HMGCR and LDLR gene expression levels as compared to wt as well, indicating that the little CETP expressed on a standard diet already redistributed cholesterol (**Figure 2 B, C**). Additionally, we assessed two Alzheimer’s risk genes, the ATP-binding cassette transporter A7 (ABCA7), a lipid transporter that is also regulated by SREBP-2, and triggering receptor expressed in myeloid cells 2 (TREM2), a lipoprotein receptor (31–34). Protein levels of ABCA7 were increased 2-fold between wt and CETPtg on either diet and the cholesterol diet doubled expression levels (thus a 4-fold difference between the extremes, wt on standard diet compared to CETPtg on cholesterol diet) (**Figure 2E-G**). TREM2 gene transcription was also increased by both the cholesterol diet and CETP expression leading to an 8-fold increase of transcript levels comparing the two extremes (wt mice on standard diet with CETPtg mice on cholesterol diet) (**Figure 2H**). However, this transcript increase could not be replicated at the protein level, which could be attributed to overall low signal intensities in the western blot (**Figure 2E, I**).

**Figure 2:**
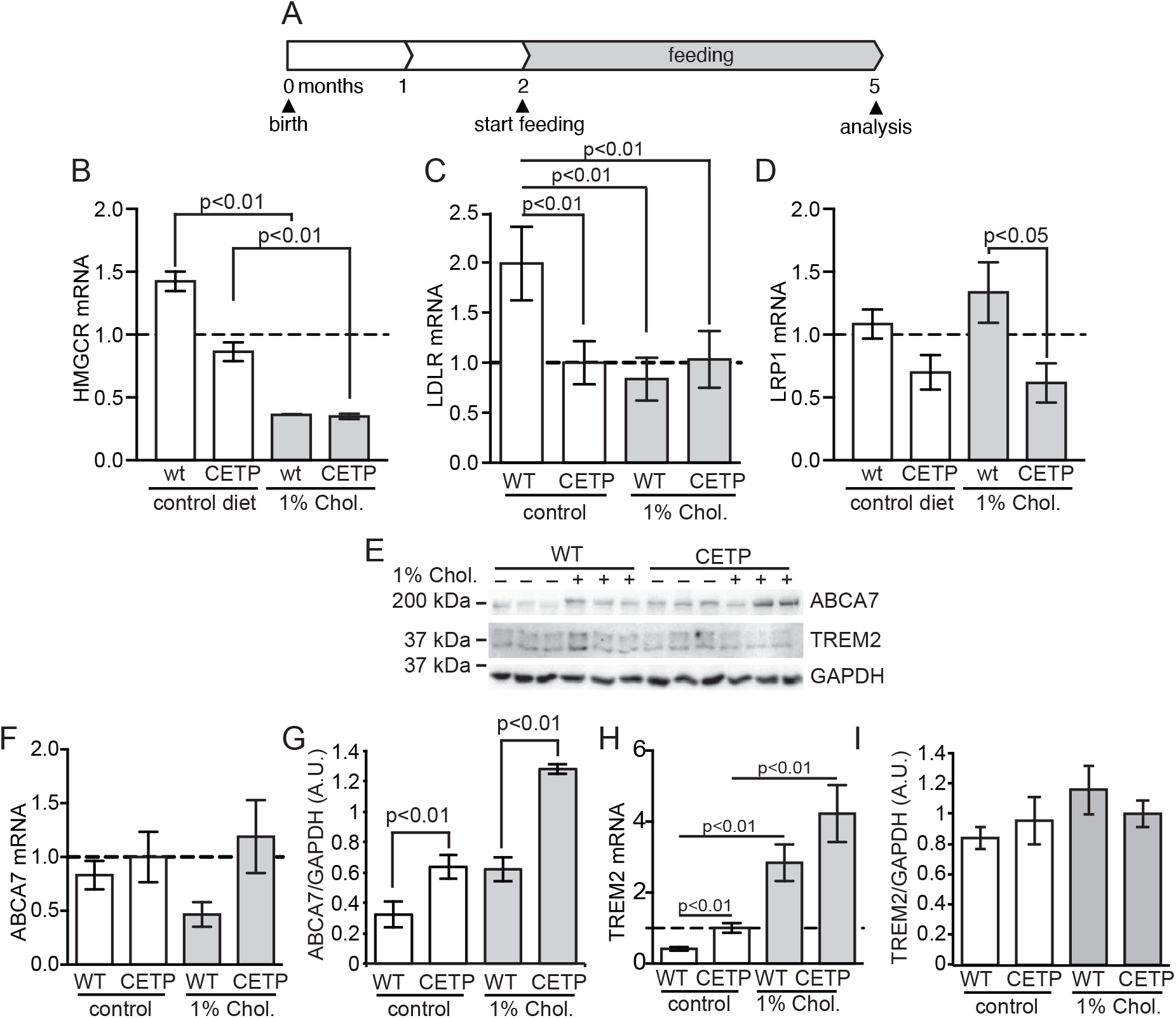
CETP promotes ABCA7 and TREM2 expression in the liver: **A:** Feeding schedule & study design. Wt and CETPtg mice were fed for 3 months starting at the age of 2 months. Biochemical analyses were performed at 5 months. **B-D:** Normalized RT-qPCR from mouse liver tissue. **B:** HMGCR, **C:** LDLR and **D:** LRP1 expression, n=6-14, mean ± SEM; 2-way ANOVA, Tukey’s multiple comparison. **E:** Western blot analysis of ABCA7 and TREM2 from liver lysates. Antibodies used: rabbit-anti-GAPDH (14C10, Cell Signaling), anti-TREM2 (Mab1729 R&D systems) and anti-ABCA7 (polyclonal, Thermo Fisher), n=6; mean ± SEM, Students T-test. **F-I:** Expression analysis of ABCA7 and TREM2 from liver samples. Normalized RT-qPCR levels of liver **F**: ABCA7 and **H:** TREM2; n=6-14, mean ± SEM; 2-way ANOVA, Tukey’s multiple comparison. Western blot quantification of **G:** ABCA7 and **I:** TREM2; n=6; mean ± SEM, Students T-test.

### CETP activity promotes peripheral inflammation

It has been previously shown that cholesterol-enriched diets induce inflammation (35). Here we assessed the effect of CETP as well as high cholesterol diet on peripheral inflammation. Specifically, we quantified the inflammatory cytokines IL1β and TNFα in mouse plasma samples using multiplex ELISA of 5 months old mice after 3 months of a 1% cholesterol or control diet. TNFα levels were significantly increased in CETPtg mice as compared to wt mice on cholesterol diet, however, it should be noted that out of the 10 plasma samples analysed, 6 samples had very low TNFα levels comparable to the control diets, and only 4 mice showed elevated TNFα levels (**Figure 3A**). Levels of IL1β were slightly increased in several CETPtg mice on the cholesterol diet, and one mouse had much higher levels **(Figure 3B)**. Since CETP is mainly secreted by the liver, we determined mRNA expression of such cytokines in the liver by qRT-PCR. As expected, the same mice with elevated plasma cytokine levels also had elevated TNFα and IL1β mRNA levels in liver (**Figure 3C, D**). Furthermore, mRNA expression of the toll-like receptor 4 (TLR4) as upstream regulator of TNFα and IL1β was also increased in mice with highest cytokine levels (**Figure 3E**). Similarly, transcript levels of IL6, an interleukin that was reported to induce the expression of lipid regulating proteins were high in 3 out of 12 mice (**Figure 3F**) (36). To analyse whether inflammatory cytokine production was extended to the central nervous system, transcript levels were determined from cortical samples. While we were able to demonstrate that CETP is expressed in the cortex of CETPtg mice, its expression levels were not affected by dietary cholesterol intake **(Figure 3G)**. Importantly, cytokine levels were not significantly increased in the brain at this age except for IL1β levels (**Figure 3H-J**). In summary, CETP expression and a cholesterol diet induced inflammatory responses in the periphery, as expected, with attenuated effects in the brain.

**Figure 3:**
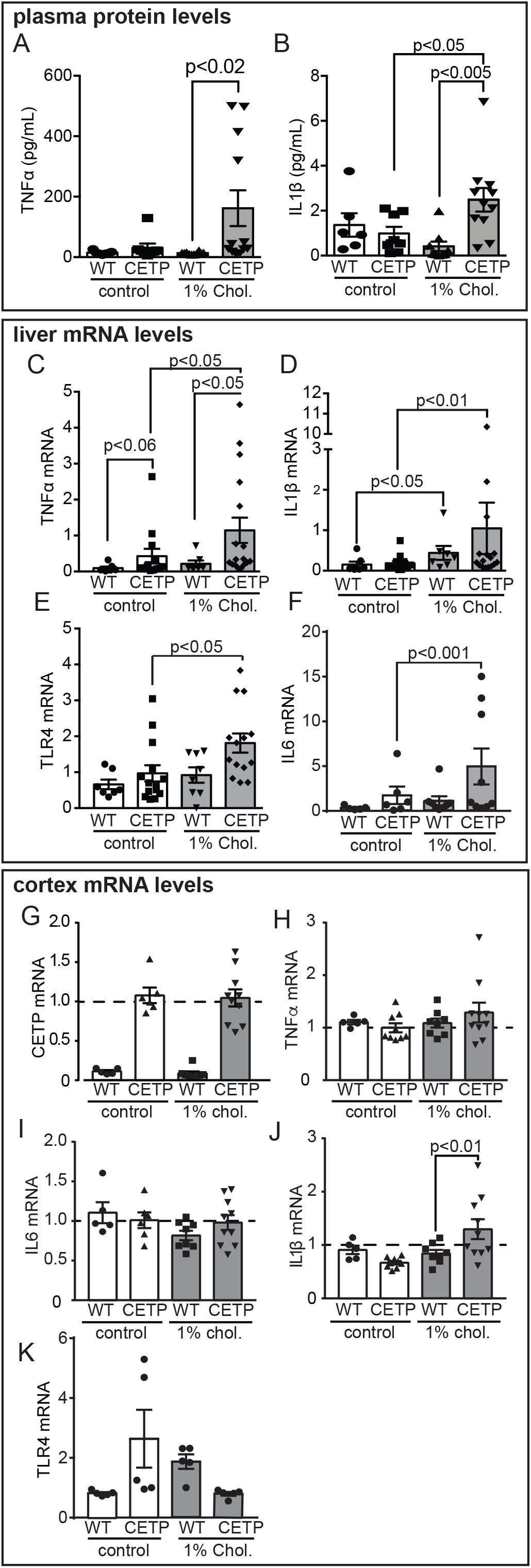
CETP activity promotes peripheral inflammation: **A, B:** Plasma cytokine levels. **A**: TNFα and **B**: IL1β measured in 25 μL plasma using a multiplex ELISA (mesoscale discoveries); n=6-11. mean ± SEM; 2-way ANOVA, Tukey’s multiple comparison. **C-F:** RT-qPCR of liver samples from 5-month old mice. Normalised expression of: **C**: TNFα, **D**: IL1β, **E:** TLR4 and **F:** IL6 expression; n=6-14, mean ± SEM; 2-way ANOVA, Tukey’s multiple comparison. **G-K:** Cytokine mRNA expression in brain samples by normalized RT-qPCR: **G:** CETP, **H:** TNFα, **I:** IL6, **J:** IL1β and **K:** TLR4 expression; n=6-10. mean ± SEM; 2-way ANOVA, Tukey’s multiple comparison.

### CETP changes the brain cholesterol composition

To determine the effect of CETP expression and high cholesterol diet on the composition and distribution of lipids in the brain, we employed matrix-assisted laser desorption/ionization imaging mass spectrometry (MALDI IMS). While several studies have looked at the distribution of lipids in the brain by IMS using 1,5-Diaminonaphthalene or other organic matrices (37,38), the visualization of cholesterol using IMS remained challenging. Here, we deposit a fine homogeneous silver layer over the tissue sections to promote the laser desorption/ionization (LDI) and allow the imaging of cholesterol and olefin containing fatty acids with high specificity and sensitivity (26). The heatmap images depict the distribution of cholesterol in sagittal mouse brain sections detected at *m/z* 493 ([M+^107^Ag]^+^ silver adduct molecular ion) **(Figure 4A)**. Cholesterol is found at the highest concentrations in the myelin-rich fibre tracts, whereas lower levels are observed in cortex, hippocampus and cerebellum **(Figure 4A**). Most interestingly, CETPtg mice showed overall higher cholesterol levels in the brain than wt mice with a 23±4% increase between wt and CETPtg mice on standard diet and a 31±4% increase between wt and CETPtg mice on cholesterol diet over the area of the whole brain (**Figure 4C, D**). The hippocampal region showed similar trends, albeit without statistically significant changes (**Figure 4E**). Since peripheral cytokine levels as well as brain IL1β mRNA levels were elevated, we further analysed levels of the fatty acid arachidonic acid as a precursor of eicosanoids and prostaglandins. Signals for arachidonic acid were comparable between genotypes and diets (while there may be a trend towards higher levels in CETPtg mice on cholesterol diet) suggesting overall low abundance of neuroinflammation in CETPtg mice at this age (**Figure 4B, F**).

**Figure 4:**
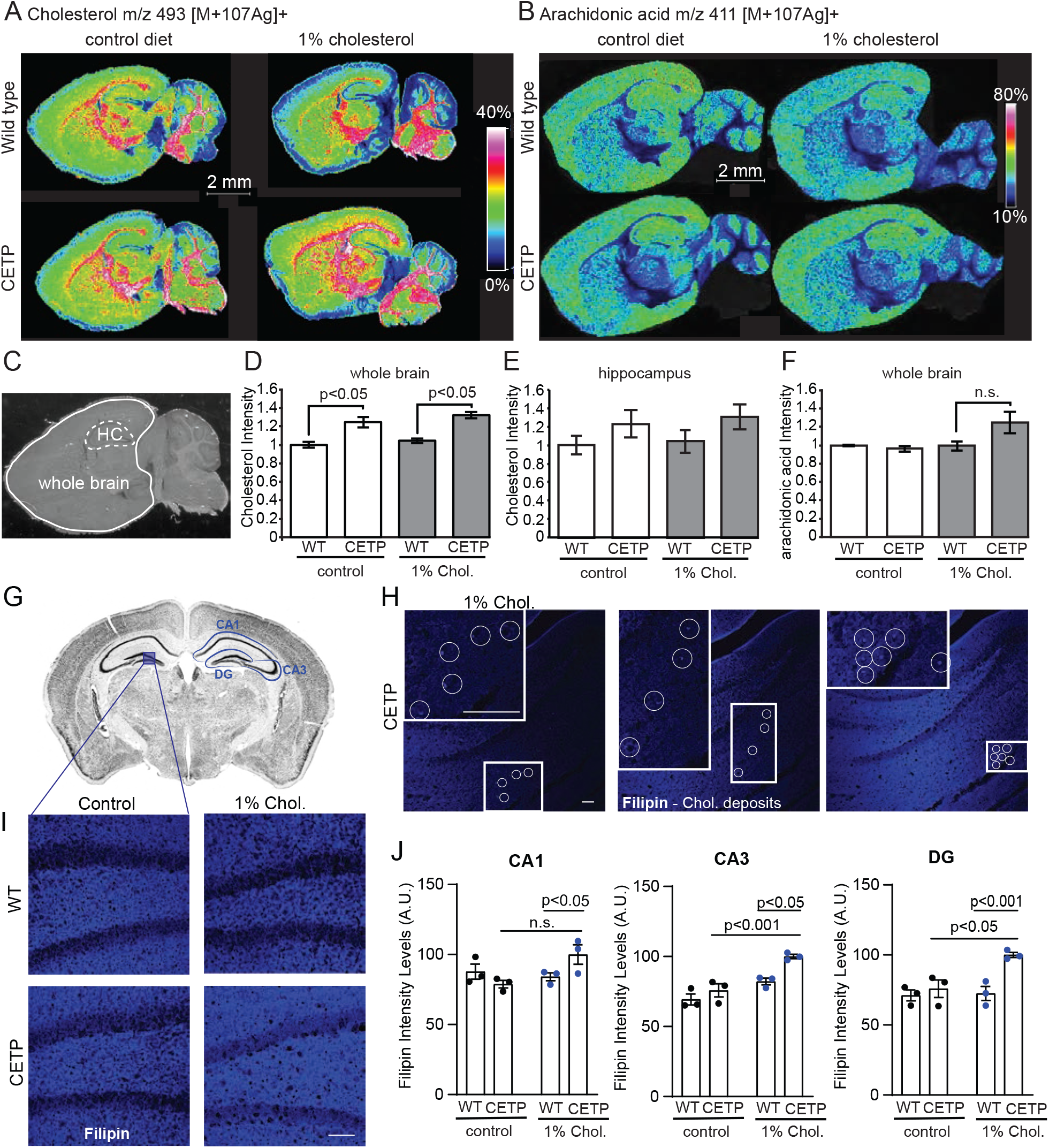
CETP changes the brain cholesterol composition: **A, B:** MALDI IMS on sagittal brain sections of 5-month-old wt and CETPtg brains. **A:** Heatmap representation of peak intensities corresponding to cholesterol (*m/z* 493 [M+107Ag]^+^) and **B** arachidonic acid (*m/z* 411 [M+107Ag]^+^). Please note the color scale with white and red depicting highest intensities. **C:** Whole brain sagittal section representing regions of interest selected for ‘whole brain’ or hippocampal (HC) quantification. **D, E:** Quantification of peak intensities corresponding to cholesterol from whole brain (**D**) and hippocampus (**E**). **F**: Quantification of peak intensities corresponding to arachidonic acid from whole brain; all n=3, mean ± SEM; Students T-test. **G:** Coronal view of mouse brain section showing hippocampal regions of CA1, CA3, and DG. The blue inset shows the portion of DG that was selected for demonstrating representative images. **H:** Consecutive confocal stacks from the hippocampus of 1% cholesterol-fed CETPtg mice showing multiple filipin-bound cholesterol deposits. Insets show enlarged views of areas with cholesterol deposits. **I:** representative images of filipin staining in brain sections of wt and CETPtg mice on standard and high cholesterol diet. **J:** Quantifications of filipin fluorescent intensities in different hippocampal regions; n=3, mean ± SEM, two-way ANOVA followed by Bonferroni’s multiple comparisons test. Scale bars=100 μm.

Based on the MALDI-IMS results, the hippocampus shows elevated cholesterol levels in CETPtg mice. The hippocampus is a well-studied brain region responsible for various critical brain functions such as memory consolidation and its alterations are commonly associated with cognitive decline. To confirm the mass spectrometry results with a second independent approach and with higher resolution, we used filipin staining to assess cholesterol levels in the CA1, CA3, and dentate gyrus (DG) regions of hippocampus (**Figure 4G**). Filipin staining in the hippocampus clearly labeled the plasma membranes in all conditions. The overall cholesterol levels assessed by intensity of filipin fluorescence was elevated by 15-25% in all hippocampal regions CETPtg mice as compared to wt mice on a high cholesterol diet (**Figure 4I-J**). Interestingly, the presence of CETP in CA3 and DG significantly increased cholesterol levels in the cholesterol-diet group as compared to CETPtg on a standard chow, suggesting location specific differences in cholesterol metabolism and/or transport (**Figure 4I-J**). Furthermore, at a high magnification, we noticed accumulated cholesterol deposits in brain sections only of CETPtg mice on the cholesterol diet (**Figure 4H**).

### Transcriptional changes in CETPtg brains induced by presenilins

To investigate whether the increased brain cholesterol levels are a result of changes in the transcription of genes that induce cholesterol synthesis, we performed a microarray from purified astrocyte mRNA (**Figure 5**). The two extreme conditions of lowest and highest cholesterol content in the brain were chosen (i.e., wt mice on a control diet compared to CETPtg mice on cholesterol diet resulting in a ~25% cholesterol increase, (**Figure 4D, J)**). Cells positive for the glutamate aspartate transporter (GLAST), a specific marker of astrocytes, were enriched from freshly dissected and dissociated whole brains using the ACSA-1 MicroBead Kit. To verify the enrichment of astrocytes, approximately 8×10^5^ cells were stained for the astrocyte marker, glial fibrillary acidic protein (GFAP), and analysed by flow cytometry revealing a purity of more than 80% across all samples (**Figure 5A**). Of note, there may be basal expression of GLAST in some neurons (39).Using total purified mRNA, CETP expression was validated in the astrocyte mRNA by qPCR (**Figure 5B**) and astrocyte transcripts were analysed on a Clariom S microarray. 595 genes were significantly up and 431 genes significantly down regulated at a threshold level of 1.5-fold change (**Figure 5C, D**). Interestingly, genes involved in cholesterol or lipid synthesis were not among the most differentially regulated genes (**Figure 5E**). In fact, such genes were downregulated such as HMGCR (1.57-fold down), SREBF1 (1.71-fold down), SREBF2 (1.84-fold down), and mevalonate kinase (MVK, 1.42-fold down). In addition, mRNA levels of LDLR and LRP1 were reduced (**Figure 5E**). Overall, this data implies that it is unlikely that increased *de novo* cholesterol synthesis is responsible for the elevated cholesterol levels in CETPtg mice.

**Figure 5:**
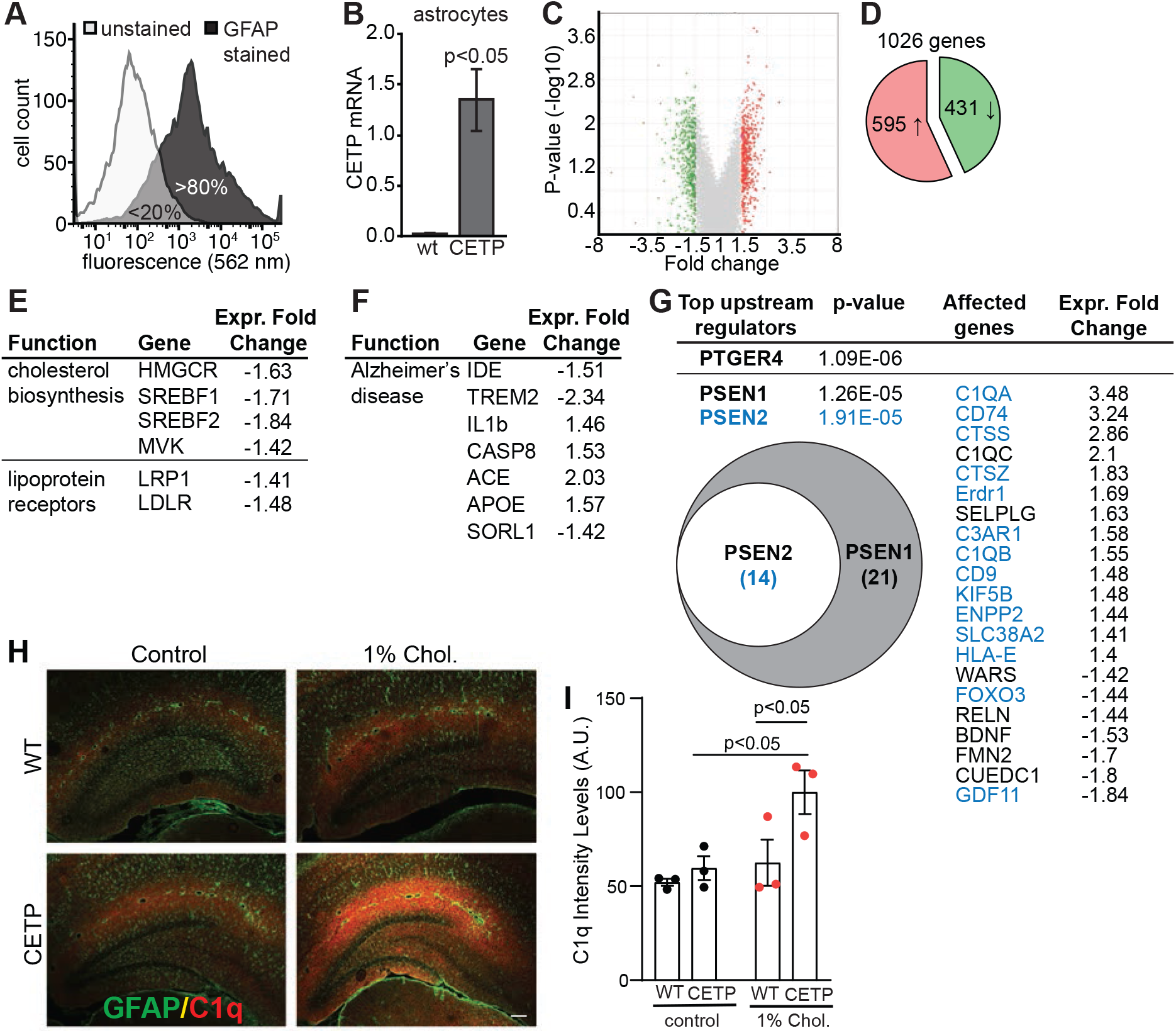
Transcriptional changes in CETPtg brain induced by presenilins: **A:** Flow cytometry analysis of astrocyte purification from 5-months-old mouse brains. GLAST-positive astrocytes were stained with GFAP. More than 80% of purified cells were GFAP positive. **B:** CETP RT-qPCR of astrocyte mRNA, n=2-3. mean ± SEM; 2-way ANOVA. **C:** Volcano plot of the mouse microarray results. Each dot represents an individual gene. The p-value of plotted against the gene regulation fold change of the corresponding gene. P-values cut-off for significance was set to <0.05. **D:** Overall, 595 genes were found to be significantly up-regulated and 431 genes were found to be significantly down-regulated in our data set. **E:** Genes involved in the *de novo* synthesis of cholesterol, generation of arachidonic acid and lipoprotein receptors. **F:** Alzheimer’s disease risk genes regulated in our data set. **G:** Pathways analysis of upstream regulators. Analyzing the fold changes in the dataset, PTGER4, presenilin 1 (PSEN1) and presenilin 2 (PSEN2) are the top 3 predicted upstream regulators. A total of 21 genes that have been reported to be regulated via PS1 and have been found in our dataset. 14 of these genes have also been reported to be regulated via PS2 (highlighted in blue). Upstream regulator analysis was performed using ingenious pathway analysis. **H:** Representative images of astrocytes (green) and C1q (red) immunostaining in brain sections of wt and CETPtg mice on normal and high cholesterol diet. **I:** Quantifications of C1q fluorescent intensities in hippocampus n=3, mean ± SEM, two-way ANOVA followed by Bonferroni’s multiple comparisons test. Scale bar=100 μm. **NCBI gene numbers:** HMGCR: 15357; SREBF1: 20787; SREBF2: 20788; MVK: 17855; LRP1: 16971; LDLR: 16835; IDE: 15925; TREM2: 83433; IL1B: 16176; CASP8: 12370; ACE: 11421; APOE: 11816; SORL1: 20660; C1QA: 20660; CD74: 16149; CTSS: 13040; C1QC: 12262; CTSZ: 64138; Erdr1: 170942; SELPLG: 20345; C3AR1: 12267; C1QB: 12260; CD9: 12527; KIF5B: 16573; ENPP2: 18606; SLC38A2: 67760; HLA-E: 15040; WARS: 22375; FOXO3: 56484; RELN: 19699; BDNF: 12064; FMN2: 54418; CUEDC1: 103841; GDF11: 14561

Since we were interested to reveal if mice with humanised cholesterol metabolism show changes in the brain relevant to Alzheimer’s disease, we analysed the effect of CETP on genes linked to Alzheimer’s (**Figure 5F**). Seven genes were identified to be differentially expressed: Upregulated genes included 1) the prime Alzheimer’s risk gene apolipoprotein E (ApoE, 1.57-fold up) involved in lipid transport, which is interesting since several epidemiological studies suggested an interaction between CETP and ApoE in the context of Alzheimer’s disease (16,18); 2) The angiotensin-converting enzyme (ACE, 2.03-fold up) producing the vasoconstrictor angiotensin II, which is upregulated and implicated in hypoperfusion in Alzheimer’s disease (40); 3) Caspase-8 (CASP8, 1.53-fold up) as a part of the apoptotic machinery, for which polymorphisms have been associated with Alzheimer’s disease (41,42); and 4) IL1β (1.46-fold up), an inflammatory cytokine that is elevated in Alzheimer’s disease brains (43). Downregulated genes included 5) the insulin-degrading enzyme (IDE, 1.51-fold down), which has been implicated in the degradation of Aβ peptides and was associated with sporadic Alzheimer’s disease (44); 6) TREM2 (2.34-fold down), which has been genetically linked to Alzheimer’s disease, acts as a lipoprotein receptor and has been intensively studied in activated microglia (33). 7) Sortilin-related receptor 1 (SORL1, 1.42-fold down), which was described to shuttle APP away from subcellular locations of Aβ production (45). Together, all these changes are in line with pathological changes in Alzheimer’s disease and imply that due to the presence of CETP several molecular changes co-occur. Next, we performed an upstream-regulator analysis, which identifies common regulators that may account for the overall changes in mRNA expression in the dataset. The top upstream regulator was the prostaglandin E2 EP4 subtype (*PTGER4*) as 24 downstream targets of *PTGER4* were differentially regulated (**Figure 5G**). *PTGER4* encodes for a G-protein coupled receptor that binds prostaglandin E2 (PGE2) and has been associated with neurotoxicity and neuroinflammation (45,46). Most interestingly, the second and third hit of upstream regulators are presenilin-1 and −2 (PSEN1 and PSEN2), the catalytic subunits of γ-secretase, an important protease in the etiology of Alzheimer’s disease, which cleaves multiple substrates and is responsible for generating Aβ peptides. Presenilin-1 and −2 were identified by 21 and 14 known downstream target genes, respectively (**Figure 5G**). To validate the expression changes found for these target genes at the protein level, we performed immunohistochemistry on hippocampal sections for one of our major hits, the initiating factor of the classical complement cascade, C1q. All genes coding for C1q protein are downstream to PSEN1 and PSEN2 and are significantly upregulated (C1qA: 3.48-fold change, C1qB: 1.55-fold change and C1qC: 2.1-fold change). Moreover, C1q is an important marker of neurodegeneration and contributes to synapse loss in Alzheimer’s disease (47,48). Immunohistochemitry for C1q protein revealed a significant increase in C1q protein expression throughout the hippocampus of high cholesterol-fed CETPtg mice compared to wt and CETPtg mice on a normal diet (**Figure 5H-I**). Overall, our results suggest that the presence of CETP and the subsequently ‘humanised’ cholesterol transport activates presenilin signaling and the complement system in the mouse brain.

### CETP activates γ-secretase

Given the elevated brain cholesterol levels and the associated stimulation of γ-secretase-mediated signalling, we investigated if CETP activity stimulates γ-secretase activity *in vitro*. To this end, we took advantage of a well-known γ-secretase substrate, notch. Once the notch intracellular domain has been released by γ-secretase, it activates transcription of notch target genes, i.e., HES1 (Hes Family BHLH transcription factor 1) and p21 (cyclin dependent kinase inhibitor 1A) (49). We therefore expressed CETP or an inactive CETP mutant (L457/M459W (50)) in HEK293T cells and determined γ-secretase activity by quantifying notch-target gene expression by qPCR (**Figure 6A, B**). Indeed, active CETP increased HES1 and p21 expression, whereas the catalytically inactive CETP had no effect (**Figure 6B**). The data shows that CETP activity causes cellular changes that stimulate γ-secretase activity *in vitro* and *in vivo.*

**Figure 6:**
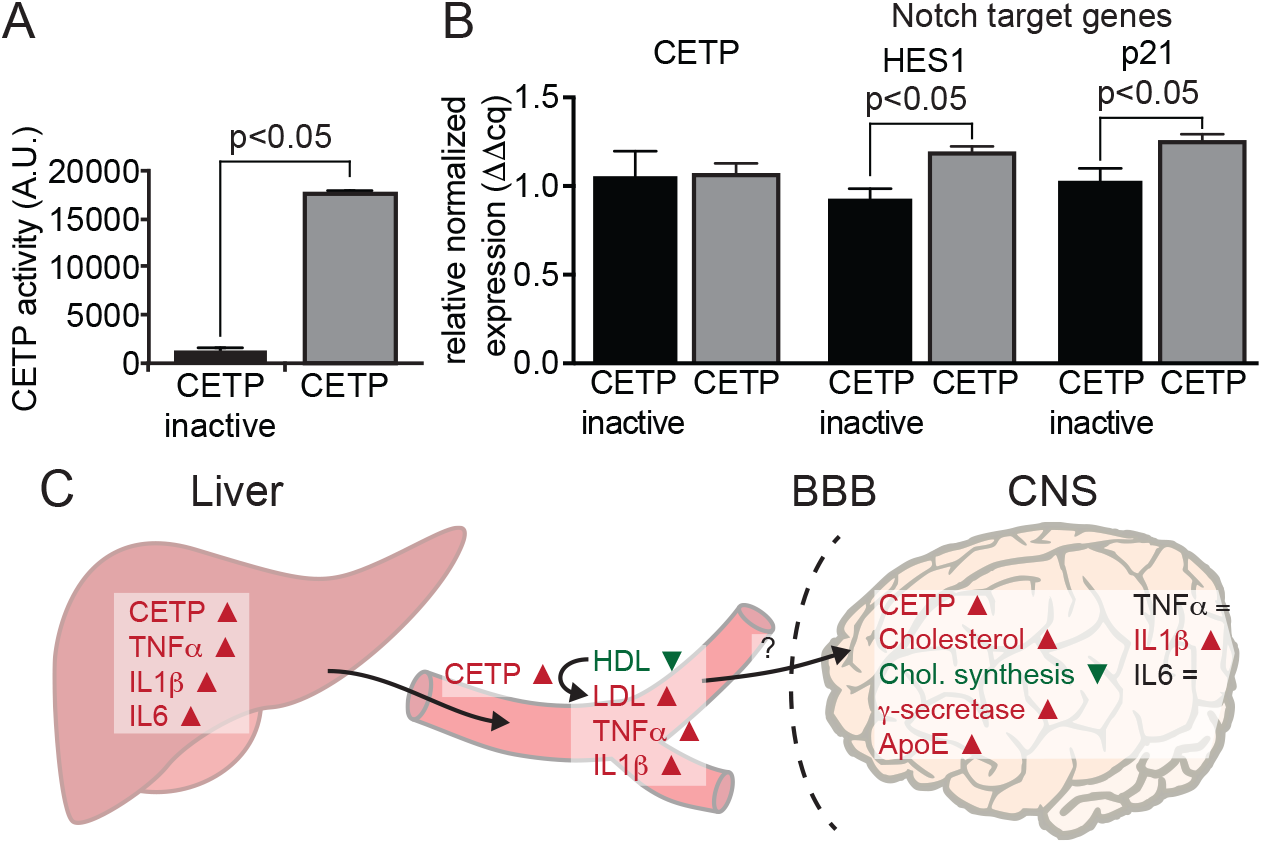
CETP activates γ-secretase. **A:** CETP activity assay of HEK293T cells transfected with wild type (wt) CETP or an inactive mutant (CETP M457/L459W). N=3, mean ± SEM, students-T test **B:** Normalised relative expression of CETP, HES1 and p21; n=3, mean ± SEM, students T-test. **C:** Schematic representation of changes observed in liver, plasma and brain in CETPtg animals as compared to wild type.

## Discussion

### CETP-mediated increase in brain cholesterol

In this study, we aimed to understand effects of CETP on brain lipid composition and gene regulation. Based on our analysis, CETPtg mice show a lipoprotein profile in the blood that resembles much better the human lipoprotein profile and importantly shows a ~25% increase in brain cholesterol levels as compared to wt. To our knowledge, this is the first report of a transgenic mouse model showing elevated cholesterol levels to this extent. Some CETPtg mice on the cholesterol diet showed peripheral inflammation, but no elevated cytokine levels in total cortical mRNA, except for IL-1β at 5 months of age and after a 3-month long 1% cholesterol diet. The enhanced inflammatory response in liver and plasma could be attributed to higher cholesterol levels in immune cells where is was already demonstrated that cholesterol augments, for instance, TLR receptor signalling, and modulates immune cells surrounding tumors (51,52). Interestingly, most of the changed gene expression in astrocytes related to inflammatory or immune-related genes. We focused on the complement factor C1q, which is elevated in CETPtg mice on a high cholesterol diet as compared to controls (**Figure 5H, I**). C1q is being increasingly discussed in the context of Alzheimer’s disease (53). C1q was originally viewed as the initiating component of the classical complement pathway. However, there is increasing evidence that suggests various complement-independent roles for C1q in innate and acquired immunity, as well as neuronal plasticity (54). As such, C1q mediates synapse pruning and it is also associated with neuroprotective effects such as the upregulation of cholesterol metabolizing genes and decreasing cellular cholesterol content (47,55). Thus, the elevated C1q levels in CETPtg mice could be a reaction to the elevated cholesterol levels aiming to eliminate excess cholesterol from the brain.

We investigated if the increase of brain cholesterol arises from *de novo* synthesis in the brain, which we could *not* confirm by transcriptome analysis. The elevated cholesterol levels may be explained by one of the following alternative pathways. While most lipoprotein particles cannot cross the blood-brain barrier, some lipid exchange between the brain and the blood can occur (2,56,57). It is well established that beneficial dietary w3-fatty acids enter the brain (58,59). In addition, 24S- and 27-hydroxysterols efficiently cross the blood-brain barrier and polymorphisms in 24S-hydroxylase were associated with Alzheimer’s Disease (2,60). Lastly, HDL particles were described to be capable of transporting cholesterol into the brain via scavenger-receptor mediated transport or transcytosis (61,62). In addition, the function of CETP in the brain remains unclear. While CETP shuttles cholesterol between HDL and VLDL in the blood, those lipoprotein particles do not exist in the brain (7). In the brain, ApoE is the predominant lipoprotein and most lipoprotein particles are HDL-like in size and decorated with ApoE or ApoJ (8,63). While a role for CETP in the brain is not clear, it is likely that it is active as a lipid transporter. However, the interaction partners may differ, and it is a possibility that CETP is involved in cholesterol redistribution between cells or acts as intracellular shuttle between organelles. Further, CETP may be involved in the storage of lipids in microglia and astrocytes. In this line, the Morton laboratory reported a role of CETP in lipid droplet formation (64,65). We observed cholesterol accumulations in CETPtg mice on a high cholesterol diet (**Figure 4H**), however, we could not determine the exact nature of such accumulations. Consequently, it is possible that lifetime exposure to CETP activity in the brain may cause an overall retention of cholesterol in the brain, leading to increased cholesterol levels observed in CETPtg mice on either diet. It will be most interesting to reveal if blood-derived CETP, centrally expressed CETP, or both are responsible for the molecular changes of the brain described herein.

In the liver of CETPtg mice, we observed an upregulation of ABCA7 and TREM2 as compared to wt mice. TREM2 mutations associate with Alzheimer’s disease, and it was thus far discussed as an immune receptor in the brain, but it also acts as a lipoprotein receptor, particularly for ApoE-containing particles (66–68). However, two recent manuscripts linked ABCA7 and TREM2 to bile acid formation in the liver (69,70). Thus, the elevated ABCA7 and TREM2 levels in CETPtg mice on cholesterol diet in the liver may reflect an increase in bile acid formation. It is tempting to speculate that ABCA7 and TREM2 may be involved in cholesterol transport or redistribution in the brain.

### Several Alzheimer-related changes are triggered in CETPtg mice

Alzheimer’s disease is the most common form of dementia and defined by the occurrence of amyloid plaques composed of Aβ peptides. Aβ peptides are generated from the amyloid precursor protein (APP) through two subsequent proteolytic cleavages. First, the ectodomain of APP is removed by β-secretase and then the membrane-bound C-terminal fragment is cleaved by γ-secretase (71–73). It is well established that higher cellular levels of cholesterol stimulate β- and γ-secretase activity (74–78). To date, the physiological function of APP as well as the trigger that leads to Aβ production remains unclear. However, it is evident that cellular pathways that stimulate Aβ production could qualify as the underlying mechanism leading to Alzheimer’s disease. The CETPtg mice analysed herein show a ‘humanised’ cholesterol metabolism but are not causatively linked to Alzheimer’s disease. Curiously, the cerebral changes that we observed (high cholesterol levels, transcriptional changes, γ-secretase activity) resemble changes that have previously been described in Alzheimer’s disease. The upstream regulator analysis of the microarray revealed the prostaglandin E_2_ receptor EP4 (gene name PTGER4) as the most significant upstream regulator (**Figure 5G**). In the brain, the EP4 receptor binds prostaglandin E2 (PGE_2_) a key inflammatory mediator in response to circulating IL1β matching our observations of EP4 as top upstream regulator (79). However, its role in mediating an inflammatory response is not completely clear as PGE_2_ can have both pro- and anti-inflammatory effects (80–83). Yet, multiple studies have linked activation of EP4 with an increase in Aβ peptides and memory loss in the context of Alzheimer’s disease (84,85). Such effects may be explained rather through stimulated γ-secretase activity than though conventional G-protein coupled receptor signalling. Hoshino *et. al* show that upon stimulation with PGE_2_, EP4 is co-internalised with γ-secretase to endosomal and lysosomal compartments where γ-secretase activity is elevated (86). Such detrimental effects were abolished in EP4 knock-out animals or through pharmacological inhibition of EP4 (84). In line with this mechanism, activation of γ-secretase activity was indeed observed as second and third top upstream regulators identified in CETPtg mice (**Figure 5G**). γ-Secretase activity is stimulated by membrane cholesterol and co-internalization with EP4 (74,87). It is important to note that while it has been well established that EP4 internalization occurs upon PGE_2_ binding, the receptor also carries a cholesterol consensus motif and directly senses changes in cellular cholesterol levels (88). Interestingly, expression of CETP in cell culture models is already sufficient to increase γ-secretase activity. Consequently, it is likely that the effects on presenilins/γ-secretase are downstream of the elevated cholesterol levels in the brain. The altered cholesterol transport, at least in the CETPtg model presented here, drives multiple molecular changes that recapitulate changes already described in Alzheimer’s disease, suggesting that a deregulation of cholesterol homeostasis may underlie Alzheimer’s disease pathology (**Figure 6C**).

Lipidomic studies have found that abnormal plasma lipid profiles, and consequently abnormal lipid biomarker panels, yield specific markers of Alzheimer’s disease (89–91). However, most animal models focus on overexpression of mutated forms of human APP, presenilin or tau, involved in a further hallmark pathology of Alzheimer’s disease. All these mouse models have low levels of circulating LDL due to the lack of CETP and therefore do not report on the impact of cholesterol transport on Alzheimer’s pathology, which may have been underestimated thus far (22). Here, we report that mice expressing human CETP exhibit elevated levels of cholesterol in the brain. In the absence of APP or presenilin overexpression, CETPtg mice show a transcriptional profile that reflects a multitude of changes previously described in the Alzheimer’s disease. Taken together, our data suggest that a mouse model expressing CETP and APP will be a valuable tool to unravel the molecular mechanisms between the peripheral and central cholesterol metabolism, ApoE and Alzheimer’s disease.

## Acknowledgements

We thank Dr. Bernard Robaire for Affymetrix software support to evaluate the microarray data, Elizabeth-Ann Kranjec for initial MALDI IMS measures, and Dr. Sandra Paschkowsky and Sasen Efrem for valuable feedback on the manuscript.

## Author contribution

FO performed and analysed all experiments presented here with the exception of the MALDI IMS data that were acquired and analyzed by EY supervised by PC, and fillipin and C1q stains that were performed by NY supervised by ARdS. LMM designed the project. FO wrote draft, LMM, NY, PC, and EY edited and revised the manuscript. All Authors approved the manuscript for publication.

